# An allometric study of the contribution of prostrate stems to belowground development of juvenile *Fagus crenata*

**DOI:** 10.1101/2024.04.29.591533

**Authors:** Yutaro Shimizu, Yudai Tondokoro, Hideyuki Ida

## Abstract

Prostrate stems represent an important morphological component of *Fagus crenata* (Siebold’s beech). This unique stem-bending growth strategy has led to the dominance of this tree species in regions with heavy snowfall along the Sea of Japan. We investigated the early-stage aboveground–belowground dynamics of *F. crenata* by applying allometric scaling theory to analyze morphological development in saplings (aged 3–20 years). Samples were collected from 25 trees in three forests in Nagano, central Japan. The scaling exponent (*b*) demonstrated an increase in the fraction of aboveground biomass (i.e., dry mass) in relation to the overall surface area (aboveground, *b* = 0.748; belowground, *b* = 0.626) and biomass (aboveground, *b* = 1.087; belowground, *b* = 0.983). These values are highly consistent with recent field observations by other researchers. Aboveground biomass growth was supported by the increasing role of prostrate stems in belowground development (*b* = 1.114). Despite its extension belowground, the growth properties of the prostrate stem may be identical to those of shoots, as both are directly influenced by nutrient sources above the germination point. Our findings highlight the significance of the prostrate stem in supporting beech survival in areas with heavy snowfall.

## Introduction

Prostrate stems are a unique morphological trait of Siebold’s beech (*Fagus crenata* Blume), which bends its trunk in response to heavy snowfall along the coasts of the Sea of Japan (Shimano 2006). Mature beech trees have relatively high flexural strength against bending stress (Miyashita et al. 2020), enabling them to deform their trunks against high snow pressure that other species fail to tolerate. In juvenile beech trees, the trunk can bend below the snow line and rise again as the snow melts the following spring. This seasonal bending causes the trees to develop a permanent curvature at the base of the trunk. Other tall deciduous tree species do not develop this bending morphology, and therefore break upon exposure to high snowfall pressure near the Sea of Japan (Homma 1997). Therefore, forest ecologists have treated prostrate stems as a potential mechanism explaining the dominance of *F. crenata* in beech forests along the Sea of Japan (Shimano 2006). Despite accumulating studies on prostrate stems in mature stands, their development in the early life stages of beech trees has not been studied extensively.

Allometric scaling theory is an emerging method in ecology that systematically describes the transformation of body or plant parts across life stages (West et al. 1997). This method permits analysis of the scaling of plant organ size (*Y*) with overall plant mass (*X*), through the formula *Y = aX*^*b*^, where *a* is a normalization constant and *b* is a scaling exponent. This theory was pioneered through the analytical and empirical demonstration of an interspecific scaling relationship of biomass partitioning into leaves, stems, and roots across nine orders of magnitude of plant mass across diverse communities (Enquist and Niklas 2002). The authors predicted and observed that shoot vs. root distribution in woody angiosperms was allometrically related, with a scaling exponent of *b* = 1.10 ± 0.019 (Enquist and Niklas 2002). Positive allometry among angiosperms suggests that trees older than one year devote more resources to aboveground growth as they mature (Poorter et al. 2012). The scaling exponent has been shown to be largely consistent across sites with differential selection for adaptations to diverse environmental conditions (e.g., site age, absolute latitude, elevation, or number of species within the community). Scaling theory is founded on the first principles of statistical mechanics and has been thoroughly tested through field observation; it complements traditional shoot:root ratios by capturing the actual functional relationships characterizing biomass allocation among organ types. The theory is based on a rigorous theoretical underpinning and provides a viable exponent baseline for the analysis of biomass partitioning trends in *F. crenata* (Marquet et al. 2014).

Kurosawa et al. (2023) applied allometric scaling to demonstrate that *F. crenata* allocates metabolic products to its roots during early growth, gradually shifting allocation towards shoots over time; when the whole plant fresh mass exceeded 0.00108 kg, the scaling exponent *b* for shoots increased from 0.09 to 1.13, whereas that for roots decreased from 2.5 to 0.825. Thus, *F. crenata* initially invests metabolic production into root growth to enhance water uptake and minimize seedling death (Kurosawa et al. 2021). As the plant matures, it shifts energy allocation to shoot growth to transition energy intake from roots to photosynthesis.

Morphological indicators of plants can also be used to analyze the unique survival strategy by which juvenile beech trees persist through the critical early growth period (Bardgett et al. 2014). In this study, we applied allometric scaling methods to analyze biomass partitioning across shoots, prostrate stems, and belowground roots in beech saplings by systematically comparing scaling exponents for juvenile *F. crenata* morphometrics based on samples collected in this study and previous field observations of juvenile beech growth.

## Methods

In October and November 2017, we collected a total of 25 *F. crenata* saplings (height, 8.5–58 cm; age, 2–17 years) from three mature beech stands in Nagano Prefecture, central Japan. The first stand is located in Karayama (36°59’N, 138°27’E, 540 m a.s.l.), with a maximum snow depth (MSD) of 304 cm, calculated as an average of measurements taken by H. Ida from 2004 to 2017; the second stand is located on Mt. Hijiri (36°29’N, 138°02’E, 1180 m a.s.l.), with an MSD of 86 cm; and the third stand is in Ohbora (36°30’N, 138°19’E, 1253 m a.s.l.), with an MSD of 59 cm. We collected 15 saplings in Karayama and 5 saplings each on Mt. Hijiri and in Ohbora. Due to low sapling density and isolation, we refrained from extensive sampling in the latter two stands for conservation reasons. We selected the Mt. Hijiri and Mt. Karayama stands because they are representative of haplotype B populations, which receive heavy snowfall near the Sea of Japan, whereas the Ohbora stand belongs to haplotype F, which inhabits low-snowfall regions near the Pacific Ocean (Koyama et al. 2012). Initially, we aimed to consider the effects of genetic differences on divergent morphological adaptation by *F. crenata* to different maximum snow depths. However, due to our limited sample sizes, data collected from all sites were pooled for analysis.

We used a high-resolution digital caliper to measure the following aboveground traits during sample collection: stem length, stem height, stem inclination, and stem diameter at ground level. We also recorded the direction and angle of the terrain. Following these measurements, we excavated each sample from the ground, including the main roots. All samples were transported to the laboratory for air-drying at room temperature, and then we removed all leaves and recorded the belowground morphology and age of each sample. For each sample, laboratory measurements included that from the tip of the stem to the end of the taproot, the length of the taproot along the slope, the length of the taproot perpendicular to the slope, and the average stem diameter at the germination point, calculated as the average of length in two orthogonal directions. We determined the germination point by visually identifying the taproot thickness and shape, and the presence of fine roots. Based on measurements of the stem and root length and diameter at the origin, we computed the cylindrical surface area of the shoots, prostrate stems, and roots (Mori and Hagihara 1988).

Next, we cut the samples into three parts (shoots, prostrate stems, and roots below the germination point) and measured the dry mass of each part. Subsequently, we dried one sample from each site at 80°C for 72 hours to measure the dry mass and determine the average moisture content. Finally, we calculated the dry mass of all samples by multiplying the wet mass by the average moisture content.

In addition, we counted bud scale scars and tree rings to infer the ages of the whole tree and each organ. We estimated the age of the whole tree by counting the scars and rings at the germination point, and we measured the age of shoots with reference to the scars and rings at the ground origin. As scars were not present on prostrate stems, we inferred their ages by subtracting the ring counts between the ground origin and germination point.

We used R Studio v4.1.3 to plot the ranged major axis regression to evaluate the scaling relationships between the biomass of each plant part and the whole-plant biomass, as well as the derived surface area (Sibly et al. 2012). We also simulated coefficients (1000 iterations per value) to calculate confidence intervals to quantify the level of uncertainty for exponents and normalization constants (King et al. 2000).

## Result

All samples were collected in the early stage of transition from seedlings to mature trees, with a median age of 12 years (range, 3–20 years). Whole-plant dry mass ranged from 0.0001169 to 0.0194953 kg (*n* = 25; Table 1).

Biomass and surface area partitioning between above- and belowground plant parts are illustrated in Figure 1, and the data are presented in Table 2. The surface area scaling of both above- and belowground surface area showed negative allometric relationships with whole-plant mass (i.e., *b* < 1 ; aboveground, *b* = 0.748; belowground, *b* = 0.626), whereas biomass scaled nearly isometrically (i.e., *b* = 1; aboveground, *b* = 1.087; belowground, *b* = 0.983). For both surface area and biomass, aboveground scaling was significantly higher than belowground scaling, based on a 95% confidence interval (CI). The normalization constants for surface area and biomass were higher in magnitude for belowground plant parts than for aboveground parts.

**Figure 1.**
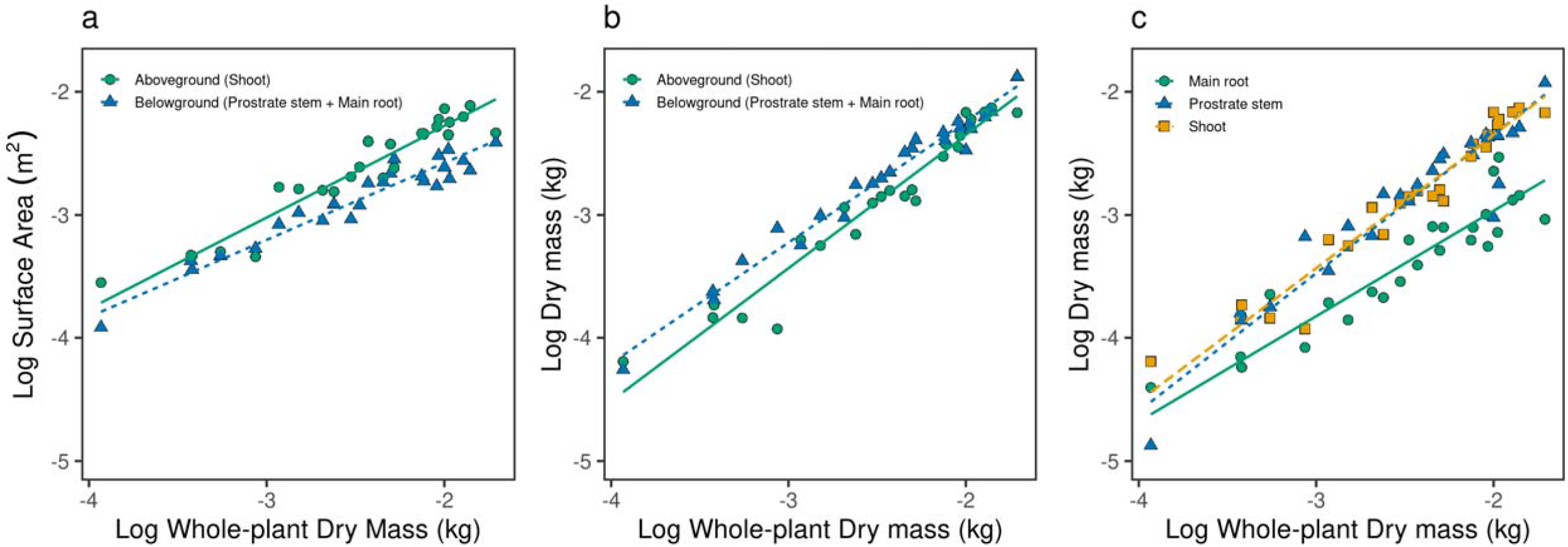
Above- and belowground partitioning of (a) surface area and (b) fresh biomass. (c) Scaling relationships among shoots, prostrate stems, and root mass in response to whole-plant fresh mass. Straight lines represent the ranged major axis regression fit for each plant part.

Prostrate stems constituted a significant portion of the belowground mass of young beech trees (Figure 1c). Compared to roots, prostrate stems alone produced a higher log (a) intercept and scaling exponent b. The biomass of shoots and prostrate stems showed consistently positive allometric scaling (shoots, *b* = 1.087, 95% CI = 0.982–1.206, *R*^2^ = 0.946; prostrate stems, *b* = 1.114, 95% CI = 0.956–1.293, *R*^2^ = 0.892), whereas that of roots showed a lower scaling exponent (*b* = 0.860, 95% CI = 0.717–1.032, *R*^2^ = 0.851). There were no significant differences in shoot and prostrate stem biomass scaling, based on the 95% CI. The normalization constants of prostrate stems and shoots were very similar (prostrate stems, *a* = -0.174; shoots, *a* = -0.126), which indicated their matched growth across whole-plant size. The scaling exponents indicated isometric scaling of shoots and prostrate stems, and that a dominant portion of the whole-plant dry mass was below the germination point (Figure 1c). The negative allometric relationship between root and whole-plant mass indicated that roots continually decrease their fraction as trees grow larger. This decreasing root fraction is compensated by consistent growth of the prostrate stem fraction. This significant scaling relationship is the main contributing factor to the apparently higher proportions of belowground surface area and biomass in young beech trees.

## Discussion

We confirmed that the prostrate stems previously reported among mature beech trees are also present in young *F. crenata* trees. When we decomposed the belowground component further into roots and prostrate stems, we found that such stems constitute a large portion of beech belowground development. The scaling exponent of these stems (*b* = 1.114) was significantly higher than that for roots (*b* = 0.860), which suggests that in addition to root growth below the germination point, submersion of the trunk within the soil is essential to the belowground support of *F. crenata* trees. Furthermore, the allometric relationship between shoots and prostrate stems showed no significant difference at the 95% confidence interval (shoots, *a* = -0.174, *b* = 1.087; prostrate stems, *a* = -0.125, *b* = 1.114). This suggests that despite their growth belowground, prostrate stems may share key developmental traits with shoots.

Kurosawa et al. (2023) reported similar results, with shoot mass exhibiting a much higher scaling exponent than root mass after a critical metabolic transition (shoots, *b* = 1.13; roots, *b* = 0.825). After consuming seed nutrients, the beech seedling first undergoes rapid root expansion to forage water and nutrients from the soil, and then develop shoots that rely primarily on photosynthesis. When the beech bends its stem into the soil, it rapidly develops adventitious roots around the ground origin to counteract additional pressure caused by deformation. This rapid growth pattern is strikingly different from that of the original roots. Similarly, our results suggest that prostrate stems maintain an identical growth pattern to shoots, which markedly differs from that of roots. The consistent positive allometric relationship detected in this study contributes a significant portion of the belowground composition of the whole plant, supporting the belowground development of young beech trees.

Young beech trees in high snowfall regions are resilient against snow pressure due to their unusually soft trunks, which are easily deformed (Meguro et al. 2015). However, as trees mature through secondary growth, they become less flexural, and prostrate stems reform within straight bark (Higuchi and Onodera 1993). Therefore, the soft bark of young trees molds an initial morphological foundation for the mature trees to also deform when exposed to high snow pressure. In addition, trees are relatively small in the early growth stage, and become larger (Weiner and Thomas 2001). Potentially, young beech trees cope with this drastic aboveground mass increase by bending growing shoots back into the soil to reinforce their physical foundation.

Given the small sample sizes that we collected in Nagano, increasing the number of samples across haplotypes would illuminate the role of prostrate stems under contrasting snow conditions. Furthermore, developing better methods to exhaustively collect the total biomass of samples is critical to the integrity of future aboveground–belowground analyses. The belowground biomass of our samples was approximately the same as that determined by Kurosawa et al. (2023), although our samples were older, which suggests that we may have missed some fine roots in our sample collection. Another important consideration is the inclusion of leaf surface area and mass. Leaves are fundamental components of allometric scaling analysis, as their photosynthetic role significantly shapes the overall growth of plants.

In summary, our findings demonstrate the important role of prostrate stems in the overall biomass and surface area of *F. crenata* during early maturity. Despite their extension underground, these stems exhibited identical growth properties to shoots, possibly because they are influenced by nutrient sources above the germination point. Our results highlight the key morphology of prostrate stems in supporting *F. crenata* survival in forest near the Sea of Japan. Elucidating the aboveground–belowground dynamics of young *F. crenata* trees will contribute to the formulation of a mechanistic understanding of the dominance of *F. crenata* in coastal regions of Japan.

## Acknowledgements

We are grateful to our colleagues from Shinshu University for their assistance with the field survey and to the staff of the Nabekura Kogen Mori-no-Ie, Iiyama City. We express our sincere gratitude to Mingzhen Lu, Leo Ware, Elianna DeSota, and Yufei Xiao for their invaluable feedback on the manuscript.

## Data availability statement

The data that support the findings of this study are openly available in Figshare at https://figshare.com/s/19ea87265ff95fb9c4cd.

## Figure and table captions

Table 1. Mass, surface area, and age profile of the collected samples.

Table 2. Scaling exponents and normalization constants for above- and belowground biomass partitioning. Belowground biomass is further partitioned into prostrate stems and roots below the germination point.

