## Supplementary material for "An allometric study of the contribution of prostrate stems to belowground development of juvenile *Fagus crenata*": Table 1

**Table 1**. Mass, surface area, and age profile of the collected samples.

| | Dry mass ($10^{-4}$ kg) | | Surface area ($10^{-4}m^2$) | | Age (years) | |
|---|---|---|---|---|---|---|
| | median | min-max | median | min-max | median | max |
| **Whole plant** | 58.082 | 1.169 – 194.953 | 52.883 | 4.037 - 100.328 | 12 | 3 - 20 |
| **Shoots** | 26.975 | 0.670 - 77.240 | 24.64 | 2.82 - 77.206 | 9 | 2-17 |
| **Prostrate stem + main roots** | 31.586 | 0.5269 – 147.810 | 18.205 | 1.217 - 39.018 | 12 | 3 - 20 |
