## Supplementary material for "An allometric study of the contribution of prostrate stems to belowground development of juvenile *Fagus crenata*": Table 2

**Table 2.** Scaling exponents and normalization constants for above- and belowground biomass partitioning. Belowground biomass is further partitioned into prostrate stems and roots below the germination point. and roots below the germination point for mass-mass scaling.

| | **Exponent (*b*)** | **95% CI of b** | **Normalisation constant (*a*)** | **95% CI of a** | ***R*** $^2$ |
|---|---|---|---|---|---|
| ***Surface area vs. Dry whole-plant mass*** | | | | | |
| Aboveground | 0.7483 | 0.6593, 0.8514 | -0.7781 | -0.9972, -0.5215 | 0.9197 |
| Belowground | 0.626 | 0.5538, 0.7068 | -1.3236 | -1.5034, -1.1225 | 0.9264 |
| ***Dry organ-specific mass vs. Dry whole-plant mass*** | | | | | |
| Aboveground | 1.0872 | 0.9815, 1.2064 | -0.174 | -0.4372, 0.1225 | 0.9461 |
| Belowground | 0.9831 | 0.9149, 1.0559 | -0.2747 | -0.4446, -0.0934 | 0.9732 |
| ***Dry organ-specific mass vs. Dry whole-plant mass*** | | | | | |
| Shoots | 1.0872 | 0.9815, 1.2064 | -0.174 | -0.4372, 0.1225 | 0.9461 |
| Prostrate stem | 1.1142 | 0.956, 1.293 | -0.1259 | -0.5189, -0.3193 | 0.892 |
| Roots | 0.8599 | 0.7170, 1.0319 | -1.245 | -1.601,-0.8168 | 0.8506 |

Note: All regressions are computed by the RMA method and $p < 0.001$. CI, confidcne interval.
